# Smoking induces coordinated DNA methylation and gene expression changes in adipose tissue with consequences for metabolic health

**DOI:** 10.1101/353581

**Authors:** Pei-Chien Tsai, Craig A Glastonbury, Melissa N Eliot, Sailalitha Bollepalli, Idil Yet, Juan E Castillo-Fernandez, Elena Carnero-Montoro, Thomas Hardiman, Tiphaine C Martin, Alice Vickers, Massimo Mangino, Kirsten Ward, Kirsi H Pietiläinen, Panos Deloukas, Tim D Spector, Ana ViñuelaX, Eric B Loucks, Miina Ollikainen, Karl T Kelsey, Kerrin S Small, Jordana T Bell

## Abstract

Tobacco smoking is a risk factor for multiple diseases, including cardiovascular disease and diabetes. Many smoking-associated signals have been detected in the blood methylome, but the extent to which these changes are widespread to metabolically relevant tissues, and impact gene expression or cardio-metabolic health, remains unclear.

We investigated smoking-associated DNA methylation and gene expression variation in adipose tissue from 542 healthy female twins with available well-characterized cardio-metabolic phenotype profiles. We identified 42 smoking-methylation and 42 smoking-expression signals, where five genes (*AHRR*, *CYP1A1*, *CYP1B1*, *CYTL1*, *F2RL3*) were both hypo-methylated and up-regulated in smokers. We replicated and validated a proportion of the signals in blood, adipose, skin, and lung tissue datasets, identifying tissue-shared effects. Smoking leaves systemic imprints on DNA methylation after smoking cessation, with stronger but shorter-lived effects on gene expression. We tested for associations between the observed smoking signals and several adiposity phenotypes that constitute cardio-metabolic disease risk. Visceral fat and android/gynoid ratio were associated with methylation at smoking-markers with functional impacts on expression, such as *CYP1A1*, and in signals shared across tissues, such as *NOTCH1*. At smoking-signals *BHLHE40* and *AHRR* DNA methylation and gene expression levels in current smokers were predictive of future gain in visceral fat upon smoking cessation.

Our results provide the first comprehensive characterization of coordinated DNA methylation and gene expression markers of smoking in adipose tissue, a subset of which link to human cardio-metabolic health and may give insights into the wide ranging risk effects of smoking across the body.

**Author Summary:** Tobacco smoking is the strongest environmental risk factor for human disease. Here, we investigate how smoking systemically changes methylome and transcriptome signatures in multiple tissues in the human body. We observe strong and coordinated epigenetic and gene expression changes in adipose tissue, some of which are mirrored in blood, skin, and lung tissue. Smoking leaves a strong short-lived impact on gene expression levels, while methylation changes are long-lasting after smoking cessation. We investigated if these changes observed in a metabolically-relevant (adipose) tissue had impacts on human disease, and observed strong associations with cardio-metabolic disease traits. Some of the smoking signals could predict future gain in obesity and cardio-metabolic disease risk in current smokers who subsequently go on to quit smoking. Our results provide novel insights into understanding the widespread health consequence of smoking outside the lung.

## Introduction

Tobacco smoking is a major environmental risk factor that predisposes an individual to chronic disease, cancer, and premature death (1, 2). Smoking directly affects exposed regions of the lung, causes damage in organs throughout the body, and results in DNA mutations that have been linked to cancer (3). The risk effects of smoking extend to multiple diseases, including cardiovascular and metabolic disease. Smoking cessation has also been linked to metabolic health, as it is associated with an increase in weight gain and in metabolic disease risk factors such as visceral fat (4).

Persistent smoking has lasting effects on DNA methylation and many epigenome wide association studies (EWAS) have identified and replicated smoking differentially methylated signals across populations with the majority of results in whole blood samples (5–19), buccal cells (20), and lung tissue (21, 22). Most smoking methylation signals show lower levels of DNA methylation in smokers and variable dynamics upon cessation. Although some alterations persist over decades, smoking cessation can result in methylation levels reverting to those observed in non-smokers, where ex-smokers exhibit intermediate methylation levels between non-smokers and current-smokers (12, 15, 17, 23). Methylation levels correlate with the cumulative dose of smoking and are associated with time since smoking cessation (12, 15, 23, 24).

Smoking can also affect gene expression, for example as reported in human airway epithelium(25, 26), lung tissue (27), alveolar macrophages (28), and lung cancer tissue (29). However, few studies have examined DNA methylation and gene expression changes concurrently, and these studies were either conducted with low coverage genome assays (such as pyrosequencing (29) and HELP assay (7)) or targeted single genes of interest in small samples sizes (7, 29).

Here we performed the first combined genome-wide analysis of smoking-related methylation and gene expression changes across tissues, focusing on adipose tissue. We identify multiple genes that exhibit both methylation and expression changes within and across tissues, showing that smoking leaves a systemic imprint on epigenetic and expression variation in the human body. Our data suggest that smoking leaves a stronger impact on gene expression, while DNA methylation smoking changes are more stable over time. By linking our findings to key human phenotypes related to cardio-metabolic health, we identify several signals that could explain some of the widespread health consequences of smoking outside the lung.

## Results

### Integrated DNA methylation and gene expression analyses in adipose tissue

Our study design is summarized in **Figure 1**. Both DNA methylation and gene expression profiles were explored in adipose tissue biopsies from 542 subjects, comprising 54 current smokers, 197 ex-smokers, and 291 never smokers. DNA methylation levels at 467,889 CpG sites from the Illumina Infinium HumanMethylation450 BeadChip were first compared between current smokers and never smokers. At a false discovery rate of 1% (P < 8.37 × 10^−7^) there were 42 differentially methylated signals (smoking-DMS) or CpG-sites, and these were located in 29 unique genomic regions comprising of 28 genes and in 1 intergenic region (**Figure 2a**). Smoking-DMS are located predominantly in the gene body (47.6%), extended promoter region (38.1%), 3’UTR (4.7%), and intergenic regions (9.5%), representing an enrichment of signals in the gene body relative to array composition. Using ENCODE ChromHMM annotations (adipose nuclei) (30), we observed that 16 smoking-DMS (38%) were located at enhancers and 9 (21%) were in or near active transcription start sites (TSS), and of these 9 were flanking bivalent enhancers (n = 3) or TSS (n = 6). As expected, methylation levels of current smokers were lower than those in non-smokers in the majority (90.5%) of the 42 signals (**Table 1**).

**Table 1.** The 42 smoking differentially methylated sites in adipose samples (smoking-DMS).

| IlmnID | CHR | Location | Gene Name | Non-smoker<br>$\beta$ (mean ± SD) | Current-smoker<br>$\beta$ (mean ± SD) | $\beta$ coef. | S.E. | P-Value | cis-meQTL | S* |
| --- | --- | --- | --- | --- | --- | --- | --- | --- | --- | --- |
| cg05951221 | 2 | 233284402 | 2q37.1 | 0.255 ± 0.054 | 0.172 ± 0.040 | -1.380 | 0.108 | 1.28$\times 10^{-29}$ | rs2853386; 3.87$\times 10^{-8}$ | |
| cg21566642 | 2 | 233284661 | 2q37.1 | 0.225 ± 0.040 | 0.167 ± 0.029 | -1.347 | 0.122 | 1.87$\times 10^{-23}$ | | |
| cg23680900 | 15 | 75017924 | CYP1A1 | 0.202 ± 0.036 | 0.155 ± 0.030 | -1.198 | 0.118 | 2.96$\times 10^{-21}$ | | O |
| cg14120703 | 9 | 139416102 | NOTCH1 | 0.748 ± 0.045 | 0.693 ± 0.044 | -1.172 | 0.118 | 1.44$\times 10^{-20}$ | | |
| cg26516004 | 15 | 75019376 | CYP1A1 | 0.696 ± 0.047 | 0.628 ± 0.058 | -1.258 | 0.126 | 1.95$\times 10^{-20}$ | | Y |
| cg10009577 | 15 | 75018150 | CYP1A1 | 0.068 ± 0.021 | 0.050 ± 0.016 | -0.810 | 0.090 | 2.48$\times 10^{-17}$ | | Y |
| cg01985595 | 6 | 136479501 | PDE7B | 0.961 ± 0.025 | 0.936 ± 0.032 | -1.015 | 0.119 | 1.09$\times 10^{-15}$ | | Y |
| cg22418620 | 5 | 172072885 | NEURL1B | 0.832 ± 0.049 | 0.765 ± 0.057 | -1.077 | 0.127 | 1.63$\times 10^{-15}$ | rs57285944; 2.15$\times 10^{-8}$ | Y |
| cg23160522 | 15 | 75015787 | CYP1A1 | 0.622 ± 0.033 | 0.583 ± 0.044 | -0.991 | 0.122 | 1.33$\times 10^{-14}$ | | Y |
| cg03636183 | 19 | 17000585 | F2RL3 | 0.506 ± 0.040 | 0.473 ± 0.038 | -0.826 | 0.103 | 1.80$\times 10^{-14}$ | | |
| cg07992500 | 2 | 37896583 | CDC42EP3 | 0.771 ± 0.051 | 0.719 ± 0.052 | -1.087 | 0.141 | 1.88$\times 10^{-13}$ | rs7595854; 1.32$\times 10^{-7}$ | |
| cg12531611 | 6 | 11212619 | NEDD9 | 0.909 ± 0.021 | 0.892 ± 0.024 | -0.855 | 0.120 | 1.12$\times 10^{-11}$ | | O |
| cg03646542 | 5 | 172076155 | NEURL1B | 0.689 ± 0.037 | 0.654 ± 0.035 | -0.880 | 0.133 | 1.87$\times 10^{-10}$ | rs7715699; 1.72$\times 10^{-10}$ | Y |
| cg00353139 | 15 | 75017914 | CYP1A1 | 0.034 ± 0.013 | 0.022 ± 0.010 | -0.787 | 0.121 | 4.47$\times 10^{-10}$ | rs11072498; 2.47$\times 10^{-6}$ | Y |
| cg21124714 | 11 | 72983097 | P2RY6 | 0.736 ± 0.037 | 0.707 ± 0.033 | -0.874 | 0.136 | 5.15$\times 10^{-10}$ | | Y |
| cg01940273 | 2 | 233284934 | 2q37.1 | 0.334 ± 0.045 | 0.302 ± 0.044 | -0.679 | 0.105 | 8.93$\times 10^{-10}$ | | |
| cg25648203 | 5 | 395444 | AHRR | 0.503 ± 0.044 | 0.459 ± 0.040 | -0.825 | 0.132 | 1.30$\times 10^{-9}$ | | |
| cg20408276 | 2 | 38300586 | CYP1B1 | 0.548 ± 0.060 | 0.499 ± 0.059 | -0.781 | 0.125 | 1.61$\times 10^{-9}$ | | O |
| cg20131897 | 12 | 52305332 | ACVRL1 | 0.694 ± 0.034 | 0.673 ± 0.028 | -0.693 | 0.116 | 5.61$\times 10^{-9}$ | rs1700159; 2.97$\times 10^{-7}$ | Y |
| cg21611682 | 11 | 68138269 | LRP5 | 0.370 ± 0.041 | 0.336 ± 0.035 | -0.734 | 0.124 | 8.10$\times 10^{-9}$ | | |
| cg19405895 | 5 | 407315 | AHRR | 0.955 ± 0.014 | 0.942 ± 0.024 | -0.768 | 0.128 | 8.38$\times 10^{-9}$ | | Y |
| cg05575921 | 5 | 373378 | AHRR | 0.713 ± 0.044 | 0.682 ± 0.039 | -0.611 | 0.104 | 1.07$\times 10^{-8}$ | rs7731963; 3.97$\times 10^{-8}$ | |
| cg13531977 | 9 | 112013420 | EPB41L4B | 0.807 ± 0.035 | 0.833 ± 0.029 | 0.831 | 0.140 | 1.14$\times 10^{-8}$ | | Y |
| cg00512031 | 4 | 5021976 | CYTL1 | 0.880 ± 0.026 | 0.855 ± 0.028 | -0.760 | 0.129 | 1.23$\times 10^{-8}$ | chr4:5022470; 1.42$\times 10^{-9}$ | Y |
| cg25189904 | 1 | 68299493 | GNG12 | 0.100 ± 0.043 | 0.064 ± 0.030 | -0.771 | 0.131 | 1.48$\times 10^{-8}$ | | |
| cg00378510 | 19 | 2291020 | LINGO3 | 0.217 ± 0.059 | 0.181 ± 0.053 | -0.781 | 0.134 | 1.53$\times 10^{-8}$ | rs12609156; 6.83$\times 10^{-18}$ | |
| cg11554391 | 5 | 321320 | AHRR | 0.065 ± 0.019 | 0.048 ± 0.014 | -0.720 | 0.125 | 2.00$\times 10^{-8}$ | | |
| cg01802380 | 13 | 107865407 | FAM155A | 0.845 ± 0.030 | 0.825 ± 0.037 | -0.737 | 0.133 | 5.69$\times 10^{-8}$ | rs9520326; 1.52$\times 10^{-12}$ | Y |
| cg14179389 | 1 | 92947961 | GFI1 | 0.083 ± 0.030 | 0.063 ± 0.028 | -0.665 | 0.122 | 1.07$\times 10^{-7}$ | | |
| cg06644428 | 2 | 233284112 | 2q37.1 | 0.036 ± 0.018 | 0.024 ± 0.010 | -0.704 | 0.130 | 1.61$\times 10^{-7}$ | | |
| cg12081267 | 2 | 98486185 | TMEM131 | 0.878 ± 0.038 | 0.858 ± 0.035 | -0.650 | 0.122 | 1.97$\times 10^{-7}$ | | Y |
| cg02162897 | 2 | 38300537 | CYP1B1 | 0.567 ± 0.060 | 0.520 ± 0.061 | -0.674 | 0.127 | 2.89$\times 10^{-7}$ | | O |
| cg11555067 | 2 | 99081350 | INPP4A | 0.725 ± 0.047 | 0.700 ± 0.046 | -0.717 | 0.138 | 3.18$\times 10^{-7}$ | rs3754893; 2.27$\times 10^{-7}$ | |
| cg04134818 | 5 | 148998446 | FLJ41603 | 0.153 ± 0.026 | 0.133 ± 0.025 | -0.690 | 0.132 | 3.26$\times 10^{-7}$ | rs11950259; 7.83$\times 10^{-6}$ | Y |
| cg03976650 | 13 | 77456505 | KCTD12 | 0.667 ± 0.061 | 0.612 ± 0.067 | -0.754 | 0.143 | 3.56$\times 10^{-7}$ | | Y |
| cg22851561 | 14 | 74214183 | C14orf43 | 0.422 ± 0.041 | 0.390 ± 0.040 | -0.634 | 0.121 | 3.92$\times 10^{-7}$ | | |
| cg10376100 | 1 | 236017278 | LYST;MIR1537 | 0.923 ± 0.036 | 0.947 ± 0.030 | 0.615 | 0.117 | 4.03$\times 10^{-7}$ | | Y |
| cg04063216 | 2 | 14772482 | FAM84A | 0.071 ± 0.016 | 0.075 ± 0.019 | 0.441 | 0.085 | 4.39$\times 10^{-7}$ | | Y |
| cg16320419 | 3 | 5025570 | BHLHE40 | 0.352 ± 0.052 | 0.315 ± 0.048 | -0.699 | 0.135 | 4.88$\times 10^{-7}$ | | |
| cg04135110 | 5 | 346695 | AHRR | 0.339 ± 0.061 | 0.384 ± 0.065 | 0.699 | 0.137 | 5.34$\times 10^{-7}$ | rs2672748; 3.42$\times 10^{-17}$ | |
| cg20109054 | 6 | 31804109 | C6orf48;SNORD52 | 0.091 ± 0.026 | 0.072 ± 0.023 | -0.659 | 0.130 | 7.85$\times 10^{-7}$ | rs3828922; 2.74$\times 10^{-5}$ | |
| cg16721845 | 11 | 68518800 | MTL5 | 0.018 ± 0.008 | 0.014 ± 0.007 | -0.530 | 0.106 | 8.37$\times 10^{-7}$ | | Y |

**Figure 1.**
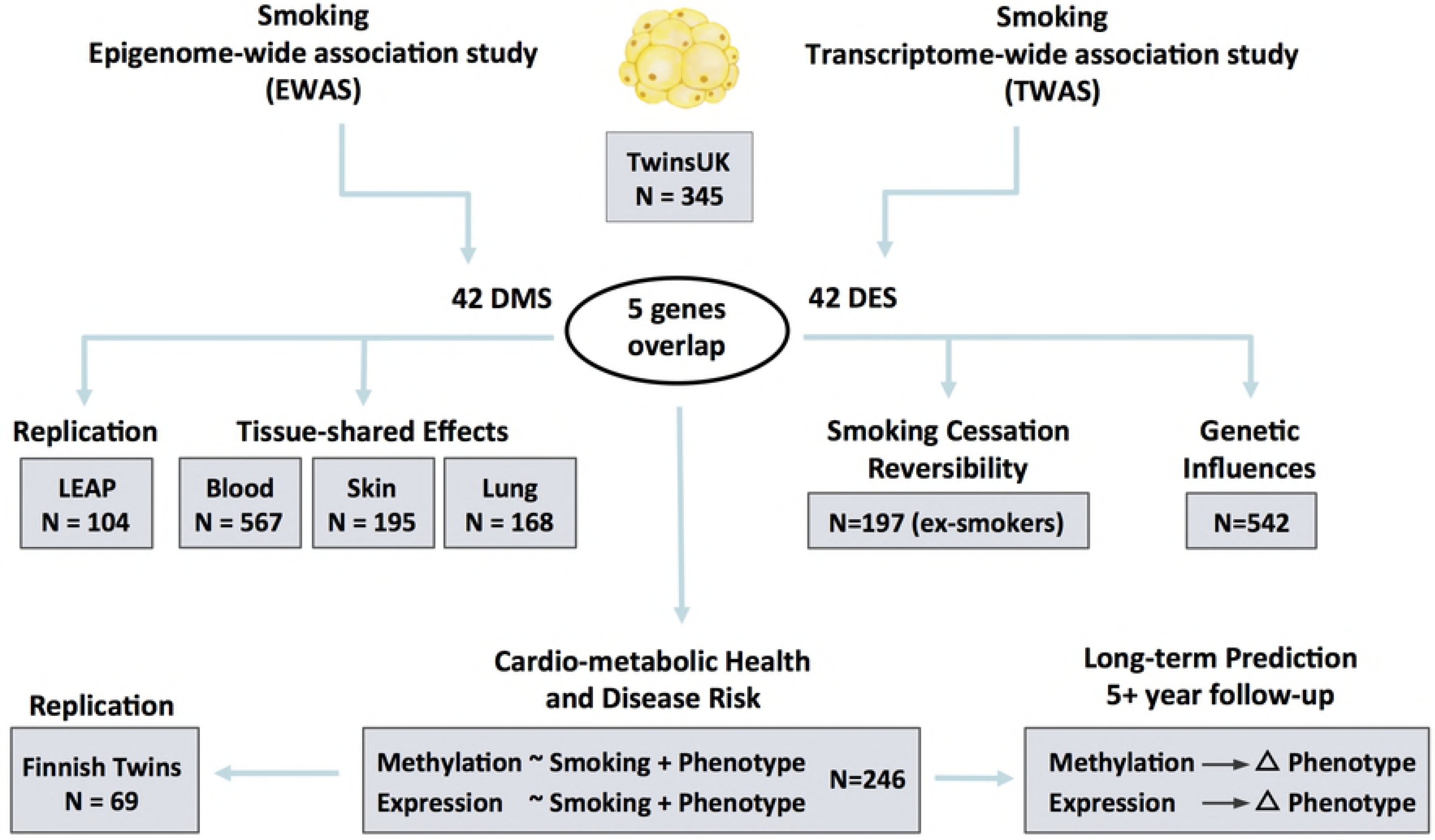
Study design. Epigenome-wide and transcriptome-wide associations studies were performed in 345 adipose tissue samples, identifying 42 smoking-DMS and 42 smoking-DES where 5 genes (14 CpG sites) overlapped. The 42 smoking-DMS were replicated in 104 independent subjects from the US, and the 14 smoking-DMS were further validated in blood, skin and lung tissue for tissue-shared effects. DNA methylation and gene expression profiles at the 42 smoking-DMS and 42 smoking-DES were tested for smoking cessation reversibility in 197 ex-smokers. Heritability and QTL analyses testing genetic and environmental influences on methylation in the 542 adipose samples were also carried out. The final set of analyses focused on exploring the link between the 42 smoking-DMS and 42 smoking-DES with cardio-metabolic phenotypes. Phenotype associations with smoking-DMS were replicated in 69 Finnish twins. The last set of analyses explored the potential of methylation and gene expression levels at smoking-DMS and smoking-DES to predict future long-term changes in adiposity phenotypes in individuals who go on to quit smoking.

**Figure 2.**
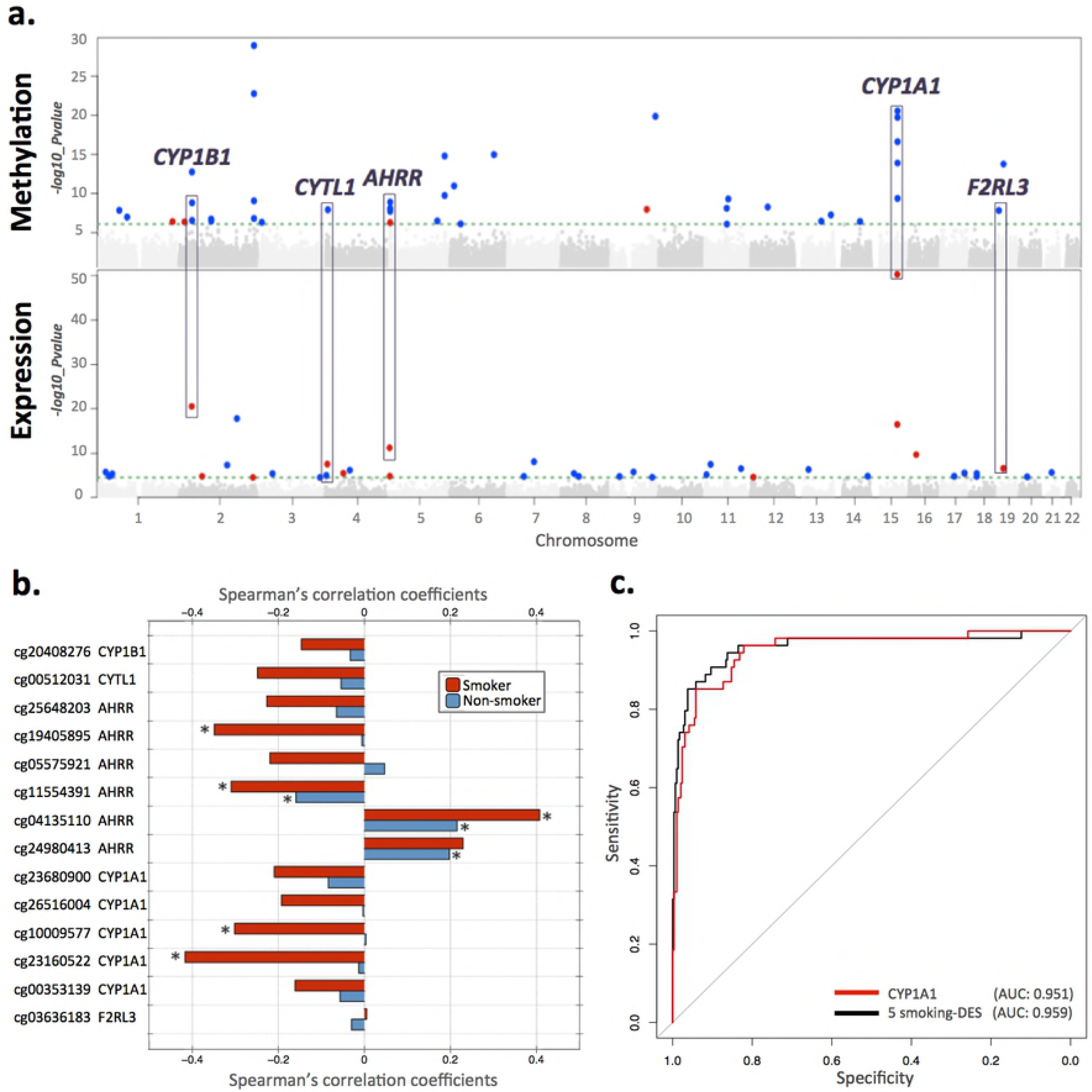
Coordinated smoking-associated DNA methylation and gene expression changes in adipose tissue. **(a)** Manhattan plots of genome-wide results for methylation (upper panel) and gene expression (lower panel) association with smoking in 345 adipose samples. Smoking-DMS and smoking-DES are indicated above the 1% FDR line (green dashed line), and are classified by direction of effect for smokers who have higher (red dots) or lower (blue dots) methylation or expression levels compared to the non-smokers. Genes highlighted by purple blocks represent 5 smoking-induced differentially methylated and expressed genes. **(b)** Methylation - expression correlation at 5 genes with coordinated smoking-DMS and smoking-DES. Pairwise Spearman’s correlation coefficients between methylation and gene expression levels for 54 smokers (red bars) and 291 non-smokers (blue bars). Asterisk indicates significance at P < 0.05. **(c)** Discrimination of current and never smokers using gene expression levels at the 5 overlapping genes. Receiver operating characteristic (ROC) curves are shown for the following combinations of predictors: *CYP1A1* gene expression level (red) and 5 smoking-DES (black) in the full dataset as an illustrative example, including AUC values from the full dataset.

To assess the impact of potential confounders on these results we performed two follow-up analyses. First we considered the impact of adipose tissue cell type composition heterogeneity, by also analyzing these data within the reference-free EWAS framework (31). We observed that the 42 smoking-DMS remained significant at FDR of 5%, suggesting that cell composition within adipose tissue did not have a major impact on our findings (**S1 Figure**). Second, habitual smoking is strongly associated with alcohol consumption (32), and in our data current smokers and ex-smokers have a higher alcohol intake compared to non-smokers (average alcohol intake = 5.96 (non-smokers), 10.03 (ex-smokers), and 11.67 (current smokers) grams per day, P = 1.06 × 10^−5^). Although our smoking analyses take into account alcohol consumption as a covariate, it is possible that the smoking-DMS in part capture alcohol consumption. To test for the co-occurrence of differentially methylated signals for smoking and alcohol consumption, we performed an alcohol-EWAS adjusting for smoking to compare the results with the 42 smoking-DMS. We observed no significant association between alcohol consumption and methylation at genome-wide significance after adjusting for smoking in adipose tissue, and only 7 smoking-DMS in *AHRR* (cg01802380, cg04134818, cg19405895), *CYP1B1* (cg19405895, cg20408276), *FAM84A* (cg04063216), and *C6orf48* (cg20109054) surpassed nominal significance (P-values between 0.05 and 0.005).

Next, RNA-sequencing profiles from the same tissue biopsy were compared between smokers and never smokers at the gene-based level using RPKM values across 17,399 genes. At an FDR of 1% (P < 2.86 × 10^−5^) there were 42 differentially expressed signals (smoking-DES) or genes (**Figure 2a**), and 14 of these were up-regulated in current smokers (**Table 2**). The most-associated expression signal was in *CYP1A1* - a lung cancer susceptibility gene, which was also one of the differentially methylated signals (**Figure 2a** and **3**).

**Table 2.**
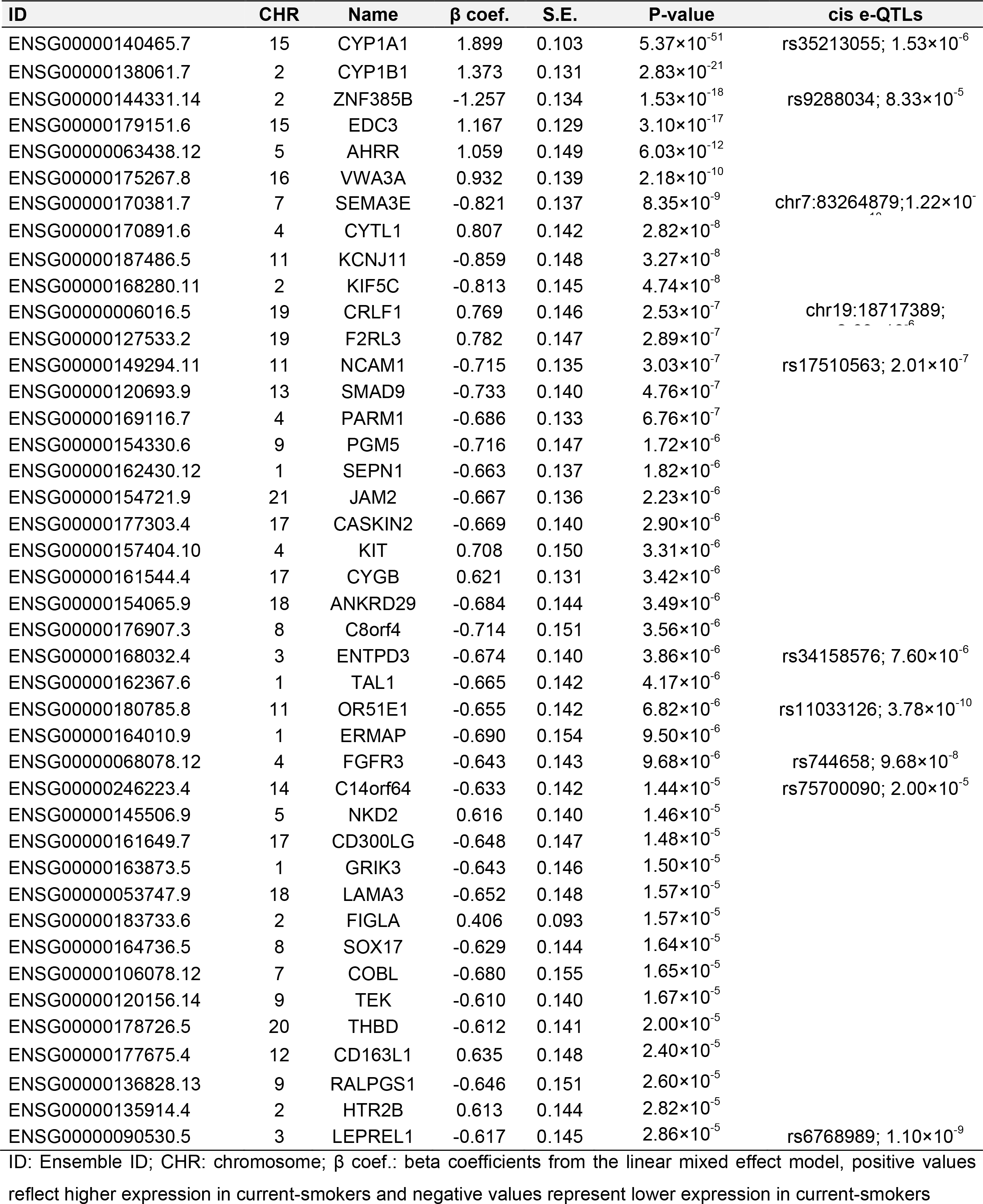
The 42 smoking differentially expressed genes in adipose samples (smoking-DES).

**Figure 3.**
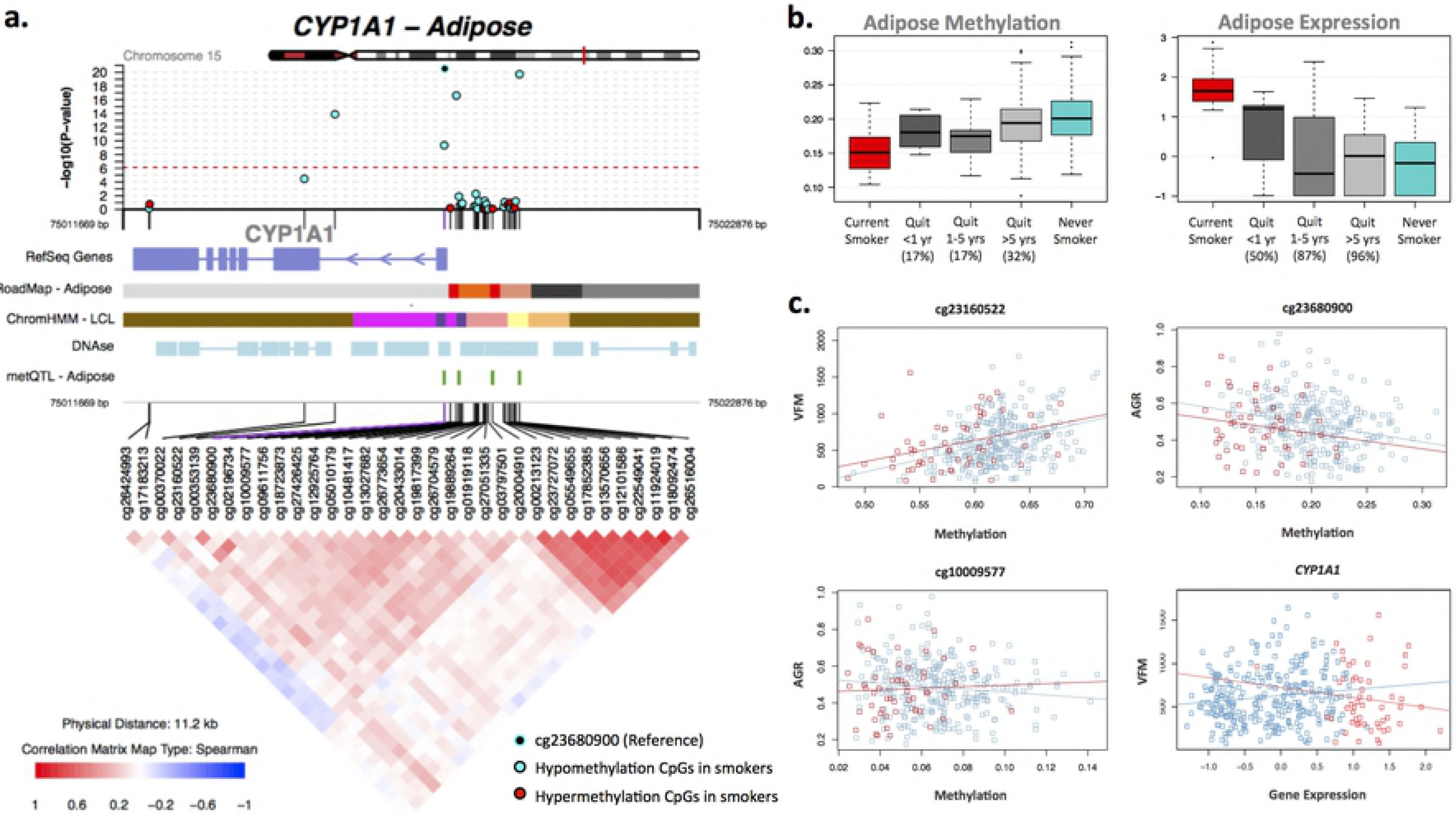
Smoking-associated DNA methylation and gene expression patterns at *CYP1A1*. **(a)** coMET plot describing the genomic region of epigenome-wide association between smoking and *CYP1A1* methylation (top panel), along functional annotation of the region (middle panel), and pattern of co-methylation at the 34 CpG sites of *CYP1A1* (bottom panel). **(b)** Coordinated DNA methylation and gene expression changes with respect to smoking cessation. Methylation (at cg23680900) and gene expression levels are shown for 5 smoking status categories: current smokers (red); subjects who quit within 1 year, subjects who quit between 1 to 5 years, and subjects who quit over 5 years at the time of methylation sampling (grey); and never smokers (blue). X-axis labels include the proportion of subjects who reverted in each smoking quit year category. **(c)** *CYP1A1* methylation associations with adiposity phenotypes, visceral fat mass (VFM) and android-to-gynoid fat ratio (AGR). DNA methylation levels at 3 CpG sites (cg23160522, cg23680900, and cg10009577 in *CYP1A1*) are shown against adiposity phenotypes in current (red) and never smokers (blue).

Comparison of the FDR 1% genome-wide significant smoking-DMS and smoking-DES showed coordinated changes at 5 genes comprising 14 CpG-sites, and these included *AHRR*, *CYP1A1*, *CYP1B1*, *CYTL1*, and *F2RL3* (**Figure 2a**). CpG-sites within *AHRR*, *CYP1B1*, and *F2RL3* were located in the gene-body, whereas CpG-sites in or near *CYP1A1* and *CYTL1* were located 200 kb to 1500 kb away from the transcription start sites. In all cases genes were up-regulated in current smokers, and in the majority of cases (93%) current smokers showed lower methylation levels compared to non-smokers. These predominantly negative correlations between methylation and expression at these five genes suggested regulatory effects (**Table 3, Figure 2b**). The methylation-expression correlations at some of these CpG sites were only observed in smokers and overall correlations were stronger in smokers compared to non-smokers.

**Table 3.** Five smoking-induced differentially methylated and expressed genes in adipose samples.

| Gene Name | IlmnID | CHR | Location | ID | r | P-value |
| --- | --- | --- | --- | --- | --- | --- |
| CYP1B1 | cg20408276 | 2 | 38300586 | ENSG00000138061.7 | -0.171 | $1.39 \times 10^{-3}$ |
| CYTL1 | cg00512031 | 4 | 5021976 | ENSG00000170891.6 | -0.176 | $1.03 \times 10^{-3}$ |
| AHRR | cg25648203 | 5 | 395444 | ENSG00000063438.12 | -0.167 | $1.80 \times 10^{-3}$ |
| AHRR | cg19405895 | 5 | 407315 | ENSG00000063438.12 | -0.134 | $1.29 \times 10^{-2}$ |
| AHRR | cg05575921 | 5 | 373378 | ENSG00000063438.12 | -0.060 | 0.2633 |
| AHRR | cg11554391 | 5 | 321320 | ENSG00000063438.12 | -0.216 | $5.37 \times 10^{-5}$ |
| AHRR | cg04135110 | 5 | 346695 | ENSG00000063438.12 | 0.279 | $1.31 \times 10^{-7}$ |
| AHRR | cg24980413 | 5 | 346987 | ENSG00000063438.12 | 0.252 | $2.10 \times 10^{-6}$ |
| CYP1A1 | cg23680900 | 15 | 75017924 | ENSG00000140465.7 | -0.329 | $3.94 \times 10^{-10}$ |
| CYP1A1 | cg26516004 | 15 | 75019376 | ENSG00000140465.7 | -0.298 | $1.70 \times 10^{-8}$ |
| CYP1A1 | cg10009577 | 15 | 75018150 | ENSG00000140465.7 | -0.266 | $5.22 \times 10^{-7}$ |
| CYP1A1 | cg23160522 | 15 | 75015787 | ENSG00000140465.7 | -0.299 | $1.48 \times 10^{-8}$ |
| CYP1A1 | cg00353139 | 15 | 75017914 | ENSG00000140465.7 | -0.222 | $3.22 \times 10^{-5}$ |
| F2RL3 | cg03636183 | 19 | 17000585 | ENSG00000127533.2 | -0.130 | 0.0159 |

IlmnID: Illumina probe ID; CHR: chromosome; Location: Illumina probe location (bp); ID: Ensemble ID; r: Spearman's correlation coefficients between methylation and gene expression data (n = 345).

To compare the impact of smoking on DNA methylation and gene expression within the same analysis framework and at a comparable scale, we used methylation and expression changes at these 5 overlapping genes (14 CpG sites) to predict a subject’s smoking status. We split the overall dataset into training and validation sets of equal size, and report here the average AUC values from 1,000 validation sets. The combination of 14 smoking-DMS levels and 5 smoking-DES levels resulted in reasonable discrimination (AUC (area under curve): 0.865). Compared to prediction results based on 14 smoking-DMS levels alone (AUC: 0.888), smoking-DES levels are better predictors (all 5 genes, AUC: 0.951). This suggests that smoking leaves a greater impact on gene expression levels, compared to DNA methylation levels at these overlapping genes. A similar high predictive value can be achieved by using gene expression levels at just a single gene, *CYP1A1* (AUC: 0.952) (**Figure 2c**). *CYP1A1* was the peak smoking differentially expressed gene, with differentially methylated signals in the promoter, and negative correlation in methylation and expression (**Figure 3a**).

### Adipose-specific and tissue-shared smoking signals

To test if the effects of smoking are shared across tissues, we first compared our adipose findings to results from whole blood samples. To this end, we tested for association between smoking and whole blood genome-wide DNA methylation (in 569 individuals) and gene expression profiles (in 237 individuals), comparing current smokers with never smokers. In blood, genome-wide significant results at FDR 1% for smoking DMS and DES overlapped at four genes (**S1 Table**). Altogether, comparison of FDR 1% significant smoking-DMS results across the adipose and whole blood datasets identified 14 CpG-sites that were genome-wide differentially methylated in both blood and adipose tissue (**Figure 4a**). The 14 tissue-shared CpG sites fell in 8 genes, including *GNG12*, *GFI1*, *AHRR*, *NOTCH1*, *LRP5*, *C14orf43*, *LINGO3*, *F2RL3*, and in the 2q37.1 intergenic region (**Table 4**). All of these sites were previously reported as smoking differentially methylated sites in blood in previous studies (5–18), and include *AHRR* - the most robustly replicated smoking-methylation signal (**Figure 5a**). DNA methylation changes in two genes (*AHRR* and *F2RL3*) that exhibit both expression and methylation smoking-associated effects in adipose tissue, were also present in blood (**Figure 4c** and **5b**).

**Table 4.**
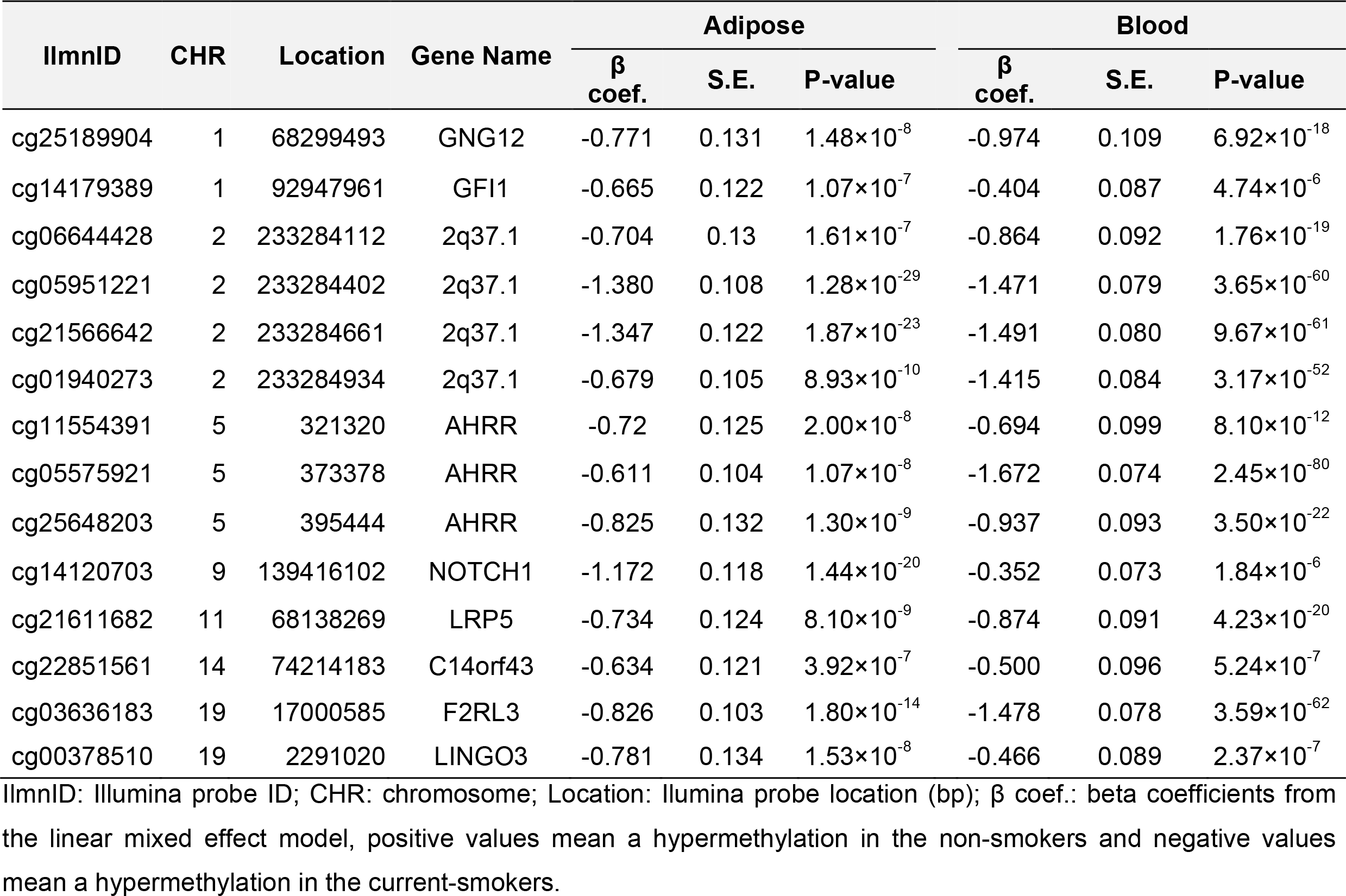
Tissue-shared smoking-induced differentially methylation sites.

**Figure 4.**
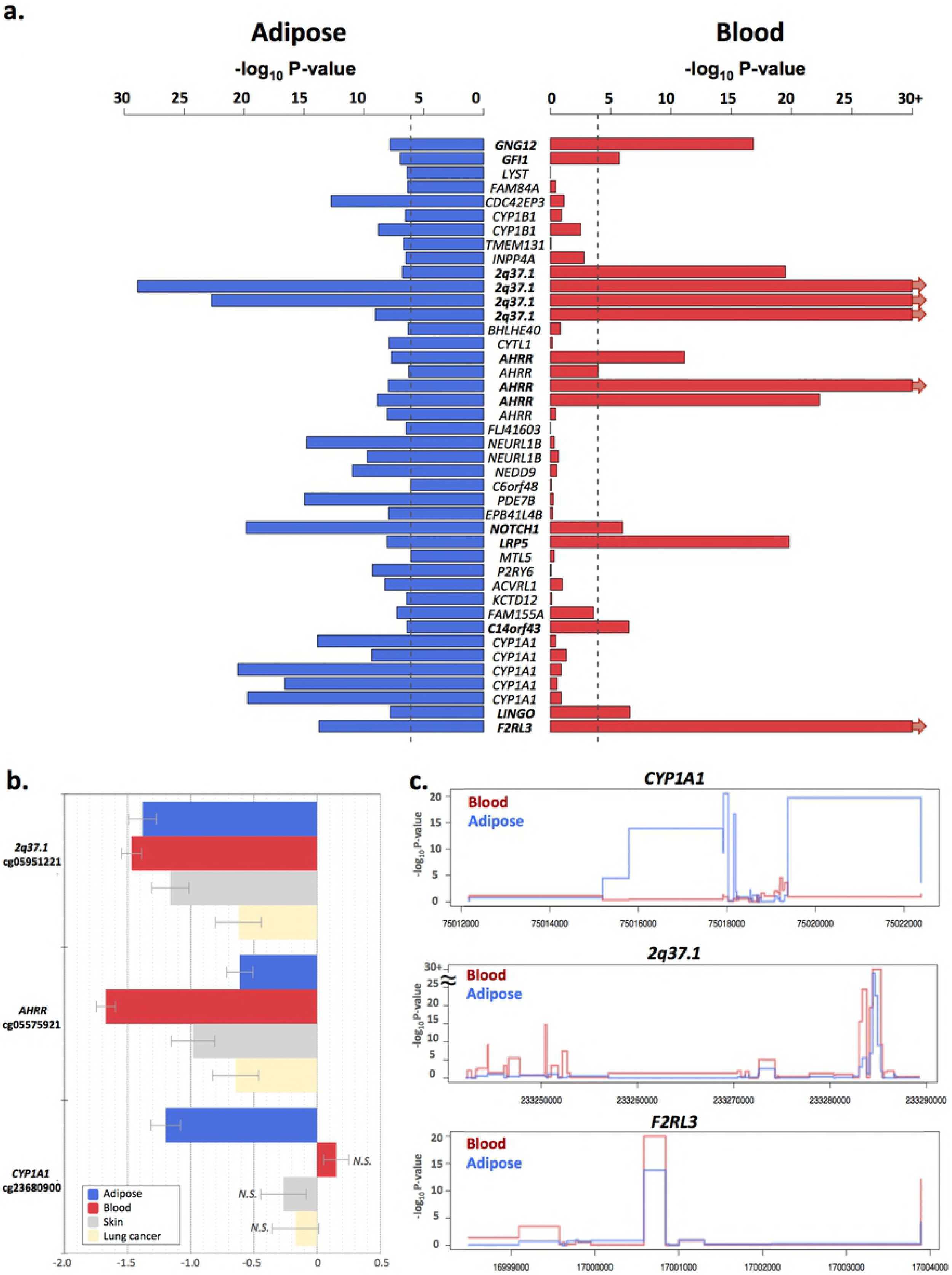
Tissue-shared and adipose-specific smoking signals. **(a)** Tissue shared DNA methylation effects across adipose tissue and whole blood. The bar-plot shows the −log_10_ P-value of the 42 smoking-DMS in adipose samples (blue), and the corresponding P-value in the blood samples (red bars). Gene names in bold denote significantly associated genes in both tissues. **(b)** Tissue-shared and tissue-specific DNA methylation effects for adipose tissue, whole blood, skin, and lung cancer tissues at 2q37.1, *AHRR*, and *CYP1A1*. Each bar represents the coefficient estimate from smoking-EWAS with standard error bars. Positive values indicate a hypermethylation in current smokers. Colors reflect tissues, with coefficients in adipose (blue), blood (red), skin (grey), and lung tissue (yellow). N.S. indicates nonsignificance. **(c)** Examples of smoking effects that are tissue-shared and tissue-specific across adipose (blue) and blood (red) samples in our datasets, including adipose-specific (*CYP1A1* in our dataset) and tissue-shared (*2q37.1* and *F2RL3*) smoking-DMS.

**Figure 5.**
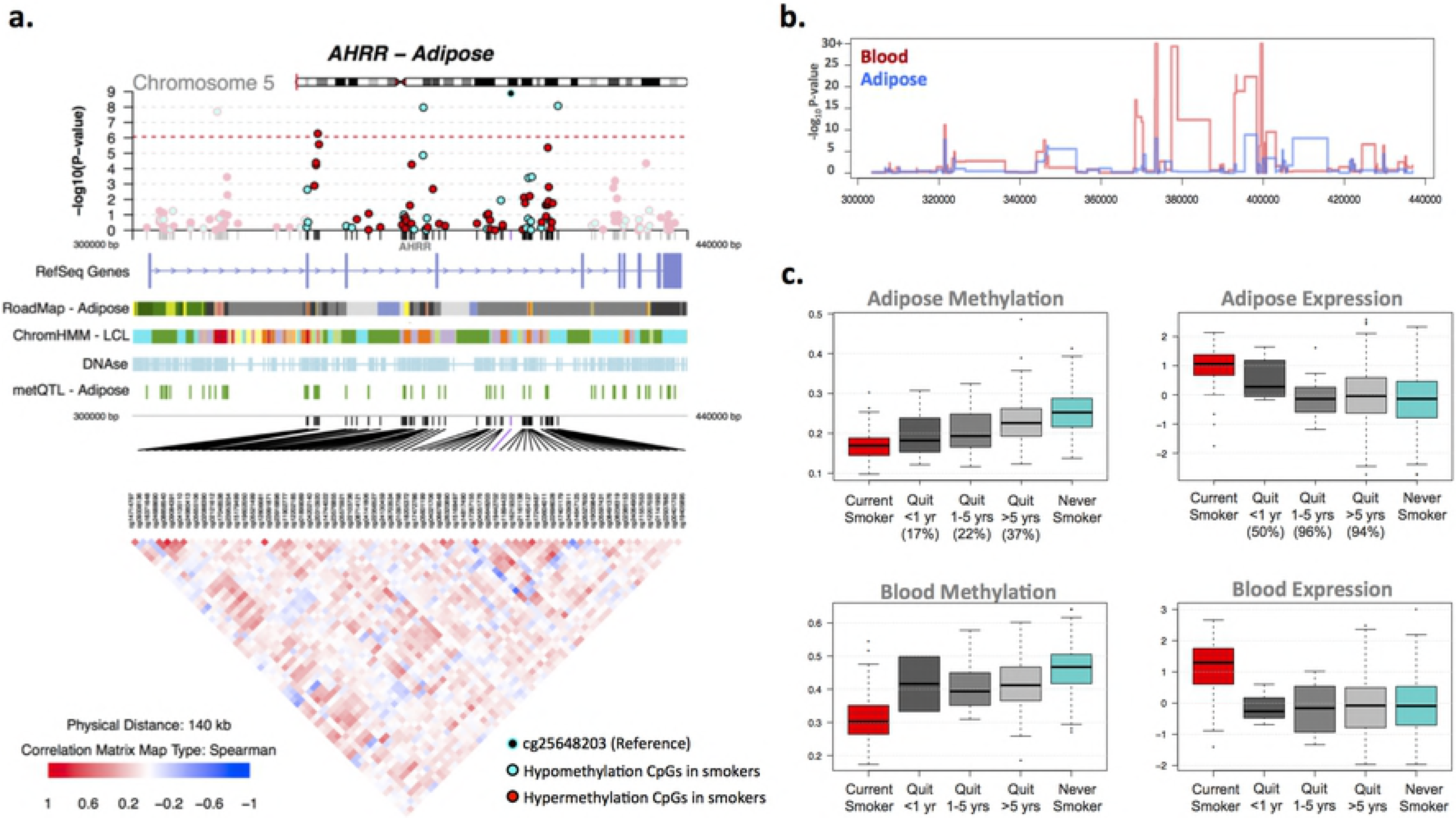
Tissue-shared smoking-associated DNA methylation and gene expression patterns at *AHRR*. **(a)** coMET plot of the association between 66 *AHRR* CpG sites and smoking. Top panel shows the −log_10_P-value of the association, the middle panel shows genomic annotation, and the lower panel shows co-methylation patterns based on Spearman correlation coefficients. (b) Tissue-shared and tissue-specific signals across CpG-sites in the *AHRR* gene region in adipose (blue) and blood samples (red). **(c)** DNA methylation and gene expression levels with respect to smoking cessation. Methylation and gene expression levels are shown for 5 different smoking status categories: current smokers (red); subjects who quit within 1 year, subjects who quit between 1 to 5 years, and subjects who quit over 5 years at the time of methylation sampling (grey); and never smokers (blue). X-axis labels include the proportion of subjects who reverted in each smoking quit year category.

We sought to validate the observed tissue-shared methylation effects at the 14 CpG-sites in additional 168 lung and 195 skin tissue samples (**S2 Table**). In lung tissue from lung cancer subjects, we validated 3 of the 14 CpG-sites in the intergenic region 2q37.1 (cg21566642 and cg05951221) and in the *AHRR* gene (cg05575921) at a Bonferroni-corrected P-value of 3.57×10^−3^. Four of the 14 CpG-sites validated in skin tissue biopsies from healthy subjects (33) in the intergenic region 2q37.1 (cg05951221, cg06644428, and cg21566642) and in *AHRR* (cg05575921). Furthermore, the majority (n = 13) of the 14 tissue-shared CpG-sites had lower methylation levels in smokers compared to non-smokers in both lung and skin methylation datasets, indicating a consistent direction of effect even if the association did not surpass significance. The smoking-DMS effect sizes observed across tissues were similar for CpG-sites in the 2q37.1 region, while the smoking effect was much greater in blood at cg05575921 in *AHRR* (see **Table 4**, **Figure 4b**).

In contrast to the methylation results, gene expression signals showed minimal evidence for tissue-shared impacts. Comparing our FDR 1% genome-wide smoking-DES across adipose and blood datasets showed that only *AHRR* was significantly up-regulated in smokers across both tissues (**Figure 5c**). *AHRR* was the only signal that showed both differential methylation and expression changes across all of the datasets that we explored in this study, including blood, adipose, skin, and lung tissue.

A proportion of our smoking-DMS and most of our smoking-DES results appear to be adipose-specific. However, the sample size of the datasets used to explore tissue-specificity in gene expression was much lower compared to that used for methylation, therefore power to detect tissue-shared effects differs across the data types. Furthermore, we are limited by access to available multi-tissue datasets for follow up, and further investigation of published findings reveals that some of our smoking adipose-specific signals have previously been detected in other tissues. For example, one of our peak results at *CYP1A1* showed methylation changes only in adipose tissue and not in blood (**Figure 4**), but has previously been reported as a smoking-methylation signal in blood (19), lung tissue (29, 34), cord blood (35), and placenta (36, 37). Unlike the persistent tissue-shared effects identified in other smoking-DMS such as signals in *AHRR* and 2q37.1, we found that smokers have lower *CYP1A1* methylation levels in adipose, skin, and lung tissue, but not in blood (19), placenta, and cord blood samples (35), overall suggesting that smoking may have contrasting effects, resulting in hyper-or hypo-methylation in different tissues (**Figure 4b**). A similar contrast in direction of smoking methylation effects is observed at smoking-DMS in *NEDD9* and *CYP1B1* across adipose tissue and in blood (**Table 1**).

### Replication of adipose smoking methylation signals

We pursued replication of the adipose-tissue smoking-DMS in an independent dataset of 104 participants from the LEAP cohort, within the New England Family Study (mean age 47 years, mean BMI 30.9, 48% male), described in detail elsewhere (38). These individuals were not affected with common diseases and had available adipose biopsy methylation profiles for 46 current smokers and 58 non-smokers. We found that the smoking-methylation direction of association was consistent at all 42 adipose smoking-DMS (**S3 Table**), and 25 of these also surpassed nominal significance in the replication dataset (P = 0.05). At a more stringent threshold the replication signal was significant at 13 sites, surpassing Bonferroni adjusted P-value for the replication analysis (P =1.19 × 10^−3^).

### Signatures of smoking cessation

We next assessed the effect of smoking cessation on the observed adipose DNA methylation and gene expression signals in ex-smokers from the discovery cohort. Here, we considered reversal for smoking methylation or expression signals to revert back to levels observed in non-smokers. We quantified the number of subjects who reverted to 25% of the change in methylation towards non-smokers, and estimated the proportion of subjects who reverted over time (in smoking-quit years), using the same approach in gene expression (see Methods).

We explored reversal patterns in adipose tissue at both the 42 smoking-DMS (**S2 Figure**) and 42 smoking-DES (**S3 Figure**), and focused on the five differentially methylated and expressed genes (14 CpG sites), where the average number of smoking-quit years was 24.8 (± 13.21) years among 190 ex-smokers. Overall, a rapid rate of reversal was observed in the first 10 years after smoking cessation, after which only subtle changes were detected in both methylation and gene-expression.

In the expression adipose data ex-smokers showed a >50% reversal rate one year after smoking cessation and reached >85% reversal after 10 years (**S3 Figure**). In comparison slower reversal was observed in the methylation dataset (**S2 Figure**). Among the 14 CpG sites only three (2 at *AHRR* and 1 at *CYP1A1*) showed a 50% reversal rate one year after cessation, while the remaining signals showed between 17% to 33% reversal (**Figure 3b** and **5c**, **S3 Figure**). Even after >40 years of smoking cessation, a proportion of smoking-DMS (n = 12; 29%) showed less than 40% reversal (**S3 Figure**). This suggests that smoking leaves a longer lasting influence on DNA methylation levels than on gene expression levels after smoking cessation.

### Controlling for genetic variation

Previous studies have shown heritable impacts on smoking behavior and nicotine addiction (39–42). We explored the impact of genetic variation on the identified smoking methylation signals. Of the 42 smoking-DMS, 14 CpG-sites had genome-wide significant meQTLs in *cis* in adipose tissue (**Table 1**). Of the 14 tissue-shared smoking-DMS, 2 in 2q37.1 and one in *LINGO3* had meQTLs in *cis* in adipose tissue, and 3 in *AHRR* and 1 in *F2RL3* had meQTLs in *cis* in blood samples.

Given our observed genetic influences on smoking-DMS, we asked if previously reported genetic variants associated with smoking behavior (41) or nicotine metabolism (42) could impact DNA methylation levels in adipose tissue. We first focused on common genetic variants that were previously associated with smoking phenotypes in the largest smoking genetic association study to date (n = 15,907) (41). We observed that all genetic variants previously strongly linked to smoking behavior (14 SNPs) (41) had an impact on adipose DNA methylation levels in *cis* (**S4 Table**). We then explored a recently reported association between a cluster of SNPs on chromosome 19 and nicotine metabolism, where the same genetic variants were also associated with whole blood DNA methylation levels in the same genomic region (42). We replicate the chromosome 19 meQTL findings in our adipose DNA methylation data in genes *CYP2A7*, *ENGL2*, and *LTBP4* (**S5 Table**), suggesting that these are strong genetic impacts on DNA methylation that are shared across tissues. Taken together, these genetic-methylation association results provide additional support for the hypothesis that some of the observed genetic impacts on smoking behavior and nicotine metabolism may be mediated by DNA methylation.

### Impacts on cardio-metabolic health and disease risk

Given the wide-ranging effects of smoking on human disease, we explored the links between the identified adipose methylation and expression smoking signals and phenotypes that are major risk factors for cardio-metabolic disease. Three metabolic disease risk phenotypes - total fat mass (TFM), visceral fat mass (VFM), and android-to-gynoid fat ratio (AGR) - were profiled using Dual X-ray absorptiometry in 288 subjects with adipose methylation and expression profiles. We assessed the association of the 42 smoking-DMS and 42 smoking-DES with these adiposity phenotypes using a two-fold approach.

First, we tested for association between adipose methylation levels at the 42 smoking-DMS and the three phenotypes, adjusting for covariates including age, BMI and smoking. We observed that smoking-DMS in *CYP1A1* and *NOTCH1* were significantly associated with measures of cardio-metabolic disease risk. Methylation levels at three CpG-sites in *CYP1A1* were significantly associated with VFM and AGR, either as main-effects (cg23160522 and VFM, beta = 1.35×10^−3^, SE = 3.03×10^−3^, P = 4.35×10^−7^; cg23680900 and AGR, beta = −1.59, SE = 0.44, P = 6.58×10^−6^) or taking into account interactions (cg10009577 and AGR, P = 5.50×10^−4^), where smokers and non-smokers have different patterns of association between DNA methylation at *CYP1A1* cg10009577 and AGR (**Figure 3c**). Probe cg10009577 is located in the *CYP1A1* promoter, suggesting gene regulatory impacts on *CYP1A1* expression levels. Correspondingly, we observed a nominally significant association between *CYP1A1* gene expression and VFM (**Figure 3c**), where smokers and nonsmokers have different patterns of association (P = 0.042). A significant negative association between DNA methylation levels and AGR was also observed with cg14120703 in *NOTCH1* (beta = −1.80, SE = 0.43, P = 1.07×10^−7^). We pursued replication of these associations in an independent sample of 69 younger Finnish twins with adipose tissue Illumina 450K methylation profiles. We replicated the overall negative association between *CYP1A1* cg10009577 and AGR (Discovery sample beta = −0.95, SE = 0.31; Replication sample beta = −0.58, SE = 0.25, P = 0.02), and observed a similar direction of interaction effects, which did not reach nominal significance in the replication sample (**S5 Table**).

We performed similar analyses with the 42 smoking-DES and observed main effects at *F2RL3* on the 3 phenotypes (VFM beta = −1.5×10^−3^, SE = 3.78×10^−4^, P = 7.8×10^−4^; AGR beta = 2.3, SE = 0.56, P = 4.5×10^−5^; TFM beta = 1.6×10^−3^, SE = 3.9×10^−4^, P = 5.8×10^−5^), and *OR51E1* on VFM (beta = −1.5×10^−3^, SE = 3.78×10^−4^, P = 7.8×10^−4^) and AGR (beta = −2.85, SE = 0.51, P = 3.1×10^−8^). We did not observe significant evidence for interaction effects in the gene expression results.

In the second set of phenotypic analyses, we explored the role of the 42 smoking-DMS and 42 smoking-DES on weight gain after smoking cessation. Recent studies have reported not only a gain in weight on smoking cessation, but also an associated increase in visceral fat (4). We considered adiposity phenotypes in 246 of the individuals in our study at two time-points, where time point 1 was the initial DNA methylation profiling and phenotype measurement, and time point 2 was a phenotype measurement on average five years later. We found that current smokers who go on to quit smoking over this five year interval show a gain in adiposity across all phenotypes (**Figure 6a**) and this effect is also observed in individuals who quit within up to four years at time point 1. However, our data suggests that this gain in adiposity is not long lasting, because we do not observe this effect in the group of ex-smokers who had quit for >5 years at time point 1. In comparison, there were no major phenotype changes within constant smokers or never-smokers across the two time points.

**Figure 6.**
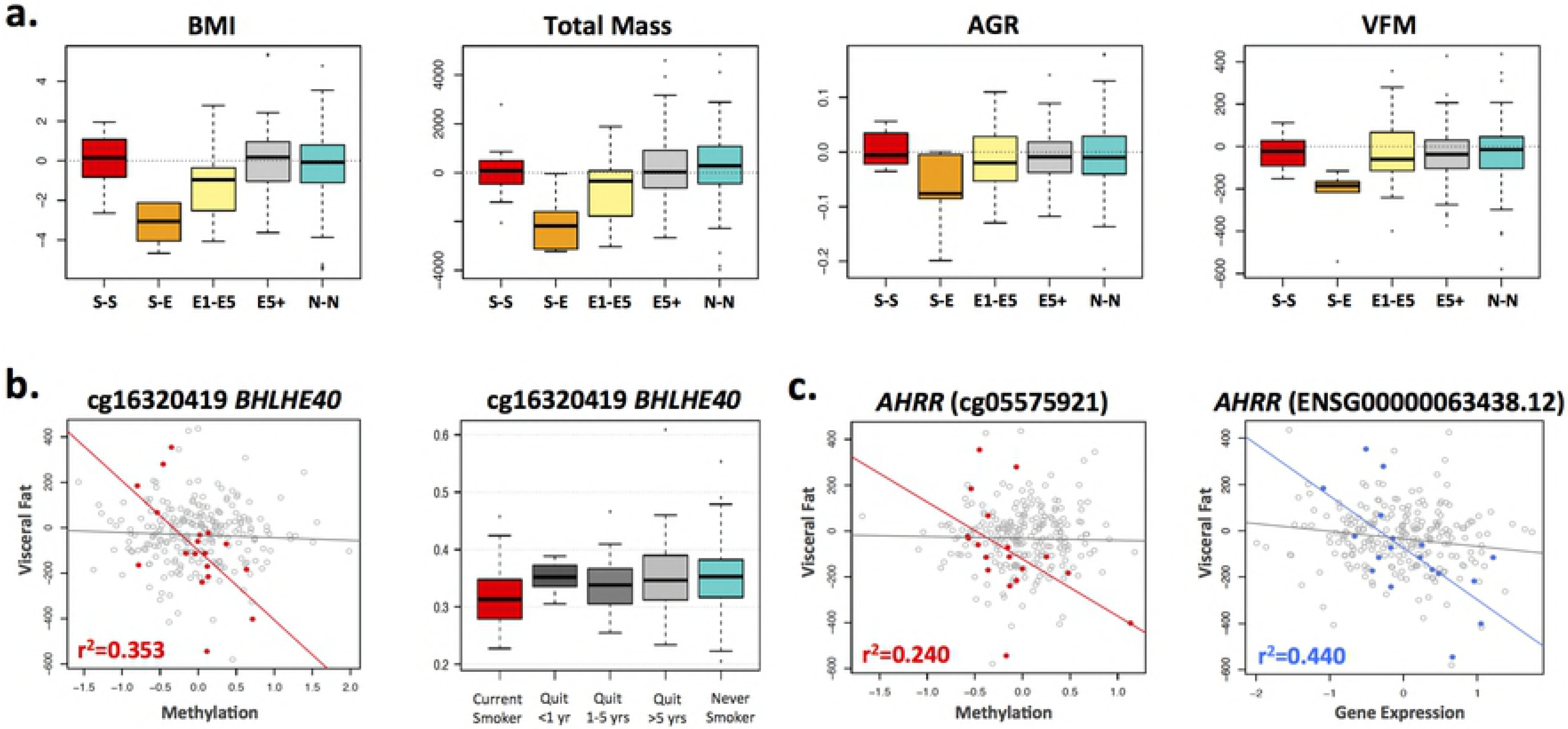
Smoking-DMS and smoking-DES relate to future changes in visceral fat mass on smoking cessation. **(a)** Adiposity phenotype changes over a 5-year time-period between time point 1 (2007-2008) and time point 2 (2012-2013). Adiposity phenotypes include BMI, total fat mass (TFM), android-to-gynoid fat ratio (AGR), and visceral fat mass (VFM). Phenotype changes are shown for 5 categories of subjects: current smokers at the two time points (S-S, n = 12), current smokers at time point 1 who quit smoking by time point 2 (S-E, n = 5), former smokers (who quit smoking within 1-5 year) at time point 1 who remain former smokers at time point 2 (E1-E5, n = 13), former smokers who quit >5 years at time point 1 who remain former smokers at time point 2 (E5+, n = 92), and non-smokers at both time points (N-N, n = 124). **(b)** Left panel shows the association between DNA methylation levels at cg16320419 in *BHLHE40* and future changes in visceral fat mass in 18 subjects in categories S-S and S-E (red points), and all remaining subjects (grey points). Right panel shows methylation cessation patterns at cg16320419 in *BHLHE40*. **(c)** Association between DNA methylation (left panel, red points) and gene expression (right panel, blue points) in *AHRR* with future changes in visceral fat mass in 18 subjects in categories S-S and S-E, and all remaining subjects (grey points).

We tested if the 42 smoking-DMS and 42 DES in adipose tissue could predict future changes in adiposity upon smoking cessation, focusing on visceral fat as the major cardio-metabolic disease risk factor. Based on the phenotype results (**Figure 6a**), we compared two groups of individuals: first, the combined group (n = 18) of current smokers at the time of methylation profiling (time-point 1) who subsequently quit smoking (n = 5), and individuals who had quit within 1-4 years at time-point 1 (n = 13); and second, the combined group (n = 228) of ex-smokers who had quit for >5 years at time point 1 (n = 92), as well as constant smokers (n = 12) and never smokers (n = 124) across the two time points. We assessed the impact of methylation or expression at the 42 smoking-DMS (**S4 Figure**) and 42 smoking-DES (**S5 Figure**) on future changes in visceral fat, selecting results that showed significantly different patterns of association in the two groups of 18 and 228 subjects.

After Bonferroni correction for multiple testing we found one DMS and one DES significantly associated with future changes in visceral fat, where a strong association effect was only observed in the group 18 subjects. This group consists of current smokers who go on to quit smoking (n = 5) and recent ex-smokers who remain ex-smokers (n = 13), and where all subjects exhibit a gain in adiposity over time. The first signal was observed in cg16320419 in *BHLHE40* (methylation by group interaction term P = 1.3×10^−4^), where methylation levels in current smokers or recent ex-smokers explain 35.5% of the variation in future gain in visceral fat (**Figure 6b**). The second signal was observed in *AHRR* (gene expression by group interaction term P = 4.7×10^−5^), where gene expression levels in current smokers or recent ex-smokers explain 44% of the variation in future gain in visceral fat (**Figure 6c**). The results were similar after correcting for smoking years and years since smoking cessation.

## Discussion

Tobacco smoking is a major disease risk factor. Our study is the first to identify smoking-associated DNA methylation and gene expression changes in adipose tissue in humans. Approximately 30% of the identified smoking-methylation signals showed significant coordinated changes in gene expression levels in 5 genes, giving insights into the cascade of molecular events that are triggered in response to smoking, toxin exposure, and nicotine metabolism. At least a third of smoking-methylation signals (in 9 genomic regions) were shared across tissues, showing that smoking leaves tissue-shared signatures. Given that our target tissue was adipose, we considered the impact of the identified smoking methylation and expression signals on cardio-vascular and metabolic disease risk. Significant associations were observed between visceral fat and android-to-gynoid fat ratio and several smoking-methylation and expression markers. Furthermore, methylation and expression levels at *BHLHE40* and *AHRR* in current smokers or recent ex-smokers were predictive of future gain in visceral fat observed after smoking cessation. Our findings provide a first comprehensive assessment of methylation and expression changes related to smoking in adipose tissue, with insights for cardio-metabolic health and disease risk.

Coordinated smoking methylation and expression changes overlapped at five genes (*AHRR*, *CYP1A1*, *CYP1B1*, *CYTL1*, and *F2RL3)*, which include well-known and strongly replicated smoking-methylation signals, such as *AHRR* and *F2RL3*. Some of these genes have previously been linked to human phenotypes. For example, GWAS associations have been reported with multiple diseases and traits, such as drinking behavior (*CYTL1*) (43), cystic fibrosis severity (*AHRR*) (44), caffeine consumption (*CYP1A1*) (45), and diastolic blood pressure (*CYP1A1*) (46); and methylation levels at *AHRR* have been linked to multiple phenotypes including lung function (47) and BMI (48). At the five overlapping genes methylation levels were predominantly negatively correlated with expression levels. CpG sites in *AHRR*, *CYP1B1*, and *F2RL3* were located on the gene-body, whereas those in *CYTL1* and *CYP1A1* were in the promoter. Our results are consistent with the expectation that promoter-based CpG-sites negatively associate with gene expression (49–51). Studies have reported both positive and negative correlations between methylation and expression for CpG-sites in the gene body (52–55). DNA methylation sites in the gene body that are negatively associated with expression levels may be located in alternative promoters that regulate the expression of particular isoforms.

*CYP1A1*, or *cytochrome P4501A1*, is a lung cancer susceptibility gene. Although in our data *CYP1A1* smoking-signals appear adipose-specific, independent studies have reported links to smoking in multiple tissues. *CYP1A1* smoking-associated methylation signals are present in lung in the fetus (56) and in adults (29, 34). In adults, effects are observed in normal lung tissue from lung cancer patients at both the *CYP1A1* promoter (34) and enhancer (29), which is also differentially methylated between normal tissue and lung tumor tissue (29). A recent large-scale metaanalysis of smoking methylation signals in blood also reported a moderate effect at *CYP1A1* (19). Maternal tobacco use was also associated with alterations in promoter methylation of placental *CYP1A1* and these changes were correlated with *CYP1A1*gene expression and fetal growth restriction (57). Furthermore, *CYP1A1* gene expression is down-regulated by *AHRR*. *CYP1A1* is inducible by agonists of the aryl hydrocarbon receptor (AhR), which include environmental pollutants and components of cigarette smoke. Following activation of AhR by an agonist in the cytoplasm, the AhR-ligand complex translocates to the nucleus, where it dimerises with the aryl hydrocarbon receptor nuclear translocator (ARNT) (58). This heterodimer binds to the xenobiotic response element (XRE) site of *CYP1A1* in the upstream enhancer region, which activates transcription. *CYP1A1* metabolizes drug molecules and environmental pollutants, including polycyclic aromatic hydrocarbons, dioxin and benzo(a)pyrene, into highly reactive intermediates. These derivatives can bind to DNA and form adducts, which may contribute to carcinogenesis (59). AhR, in complex with xenobiotic compounds and ARNT, induces *CYP1A1* expression, which subsequently detoxifies toxic components of cigarette smoke. *AHRR* suppresses *AhR* expression through binding to ARNT. Hypomethylation of *AHRR* and associated increased *AHRR* expression may therefore reduce cellular responses to smoking through *CYP1A1* (60).

In addition to *CYP1A1*, other smoking signals that we identify in this study have also been previously linked to lung cancer. *CYP1B1* differentially methylated effects have been reported for smoking, for lung cancer and for age at cancer diagnosis in nonsmall cell lung carcinoma (NSCLC) samples (61). Several of our smoking signals were previously reported to be differentially methylated in lung adenocarcinoma tumor and matched non-tumor tissue (62). These included two of our top smoking-DMS, *CYTL1* and *ACVRL1*, and seven of our top smoking-DES, *CYTL1*, *JAM2*, *CYGB*, *TAL1*, *GRIK3*, *SOX17*, and *TEK*.

In line with previous studies we observe that genetic variation can impact the smoking-DMS, with potential implications for genotype influences on the rates of toxin elimination and nicotine metabolism in the human body. Importantly, we observe that all of the major smoking genetic variants detected in the largest smoking GWAS to date appear to influence DNA methylation levels in *cis*. These findings strongly suggest that DNA methylation may mediate some of the effects of genetic influences on smoking behavior, toxin elimination, or nicotine metabolism. We also replicate results from a genome-wide association study of nicotine metabolite ratio, identifying a 4.2Mb region on chromosome 19q13 where GWAS SNPs were also associated with DNA methylation levels (42). Taken together, these findings suggests some of the observed genetic impacts on smoking behavior and nicotine metabolism may be mediated by DNA methylation, and that such effects are robust and shared across tissues.

Our analyses specifically in ex-smokers show variability in the extent of signal reversal over time, which is consistent with previous findings. We observe an overall trend towards at least partial reversal at most of the identified smoking-associated signals. Importantly, our study is the first to show that this trend is also observed in gene expression levels. Our findings suggest that smoking has a longer-lasting influence on the methylome compared to the transcriptome, where the majority of reversal effects occur within the first year after smoking cessation.

The smoking-methylation signals were assessed for association with adiposity phenotypes that constitute major cardio-metabolic disease risk. Significant associations were observed between visceral fat mass and android-to-gynoid fat ratio with methylation levels at smoking-markers with functional impacts on gene expression, such as *CYP1A1* with replication, and in signals that were shared across tissues, such as *NOTCH1*. Associations were also detected with smoking-DES. These results may help improve our understanding of how smoking impacts metabolic health, and to explore this further we considered smoking effects on future changes in metabolic phenotypes on smoking cessation. Visceral fat has a strong association with obesity-related cardio-metabolic diseases, such as type 2 diabetes and cardiovascular disease (63, 64) and is a major cardio-metabolic disease risk factor. At smoking markers *BHLHE40* and *AHRR* DNA methylation and gene expression levels in current smokers were predictive of future gain in visceral fat observed after smoking cessation. Although the sample size of current smokers who go on to quit smoking in our data is modest, these findings provide an interesting insight into potential molecular mechanisms mediating environmental effects on cardio-metabolic disease risk, and require replication in larger samples.

A limitation to our study is partial correction for the influence of expected covariates. These include first, alcohol consumption, which co-occurs with smoking. In our co-occurrence analyses, none of the alcohol-associated CpG sites reached genome-wide significance after adjusting for smoking. In a previous alcohol EWAS in blood, Liu et al. (65) also found that the effect size of the majority alcohol-DMS was not affected by smoking status suggesting that despite their co-occurrence, smoking and alcohol impact DNA methylation in different aspects. A related question is optimal correction for cell composition in adipose tissue. Since we only had access to subcutaneous adipose tissue biopsies, rather than isolated cell subtypes, we corrected for cell composition by using the analytical approach within the reference-free EWAS (31) framework and found that the majority of results remained largely unchanged. However, it is possible that this does not fully capture the effect of a heterogeneous population of cells as a confounder. Some of the smoking-DMS such as *BHLHE40*, which was also found to be predictive of future gain in visceral fat, may reflect cell-specific methylation profiles. *BHLHE40* was previously reported to be hypo-methylated in activated NK cells (but not in naive NKs, T and B-cells) (66) and a similar trend was observed for *AHRR* (66). One interpretation of these findings is that some smoking signals are cell subtype specific (67, 68), potentially reflecting a selective enhancement of activated cells, because smoking can also induce changes in blood count (69). In adipose tissue, this particular effect may be represented as an infiltration of activated NK cells, and this infiltration may increase with obesity, diabetes, and smoking. On the other hand, the relative abundance of NK DNA compared with adipose DNA in adipose tissue is minimal therefore these effects should be minimal. Future studies are needed to assess the impact of these potential confounding effects, using for example histological and immunological staining of adipose tissue.

## Conclusion

Our results show that smoking can impact DNA methylation and gene expression levels in adipose tissue. To our knowledge, this is the first study that performed genome-wide analyses of smoking in adipose tissue DNA methylation and gene expression profiles. The key results were that first, smoking leaves a signature on both the methylome and transcriptome with overlapping signals, second, smoking methylation signals tend to be tissue-shared effects, third, smoking has a longer lasting influence on DNA methylation levels than on gene expression after smoking cessation, and forth, specific smoking methylation and expression signals are associated with metabolic disease risk phenotypes as well as future weight gain after smoking cessation.

## Materials and methods

### Study population and sample collection: TwinsUK

The adipose tissue samples were obtained from 542 female twins who were recruited as part of the MuTHER study (Multiple Tissue Human Expression Resource) in TwinsUK cohort. The TwinsUK cohort was established in 1992 to recruit MZ and DZ same-sex twins (70). All twins in the current study are Caucasian females and ascertained to be free from severe disease when the samples were collected. The 542 twins included 84 MZ pairs, 112 DZ pairs, and 150 singletons. Details of biopsy procedures and sample descriptions are described previously (71). The subcutaneous adipose tissue samples for methylation and expression profiling were obtained from the same punch biopsies in the subjects’ abdominal region, and immediately stored in liquid nitrogen. Both DNA and RNA were extracted from the adipose tissue for genome-wide methylation and expression profiling. To explore tissue-shared effects, peripheral blood samples from 789 and 362 subjects for genome-wide methylation and expression profiling, respectively, were collected from twins in TwinsUK. From the 542 subjects, 200 and 222 subjects donated blood samples for methylation and expression profiling, respectively. Blood samples and adipose tissues were collected during the subject’s visit to the clinic.

### Replication and validation samples

#### Replication sample for 42 smoking-DMS: USA

The first replication sample included 104 participants from the New England Family Study, the LEAP cohort (mean age 47 years (range: 44-50), mean BMI 30.9 (range: 19.43-54.24), 48% male; see **S6 Table**), described in detail elsewhere(38). The individuals are of mixed ancestry (63.5% white) and were not affected with disease. There were 46 current smokers and 58 non-smokers. Subcutaneous adipose tissue samples in these participants were collected from the upper outer quadrant of the buttock, followed by DNA extraction, and Infinium HumanMethylation450 BeadChip array profiling as previously described(36). Replication analyses were performed using a linear regression model adjusting for age, gender, BMI, and batch effect.

#### Replication sample for cardio-metabolic health and disease risk phenotype analyses: Finland

The second replication sample included 69 Finnish twins (mean age 31 years, mean BMI 27.5, 44.9% male; see **S6 Table**), who were recruited as a part of the Finnish twin cohort. The Finnish twin cohort has been previously described in detail (72, 73). The sample included 34 full MZ twin pairs and 21 current smokers. DNA methylation profiling was measured by Infinium HymanMethylation450 BeadChip array and TFM and AGR were determined by DEXA. Replication analyses were performed using a linear mixed effect regression model adjusting for age, gender, BMI, family, batch effect, and alcohol intake. Sample characteristics of the replication cohorts are shown in **S6 Table**.

#### Validation sample for tissue-shared effects: lung tissue (74)

The first validation dataset included 168 lung cancer female subjects (mean age 65 years; see **S7 Table**), which is a subset of a multicenter cohort of 450 subjects with non-small cell lung cancer (GEO dataset: GSE39279) (74). In the validation analysis, we selected only female subjects who had smoking records (129 smokers and 39 non-smokers) and used a linear regression model to test for the effect of smoking on methylation, adjusting for age, cancer stage (1 to 4), and cancer type (adenocarcinoma or squamous). DNA methylation data were measured by Infinium HumanMethylation450 BeadChip and BMIQ normalization was performed prior to analysis.

#### Validation sample 2 for tissue-shared effect: skin tissue (33)

The second validation dataset included 195 skin tissue samples from twins (mean age 59 years; see **S7 Table**), and these subjects are part of TwinsUK. This analysis included 37 current smokers and 158 never smokers cancer-free female subjects only. The TwinsUK skin samples and the evaluation of DNA methylation in the samples are described elsewhere (33). We performed the analysis using a LME model adjusting for age, BMI, alcohol consumption, batch effect, family structure and zygosity. Sample characteristics of the two validation cohorts are shown in **S7 Table**.

### Phenotype collection

During a subject’s clinical visit, basic demographic information was collected, with onsite measurements such as height and weight, DEXA measurements, and clinical assessments. Smoking was determined from a self-reported questionnaire. There was longitudinal self-reported data on the smoking status of each subject, since twins regularly visit the research clinic. Smoking status was defined in 3 categories: current smokers, ex-smokers, and non-smokers. Current smokers were defined as those subjects who consistently smoked cigarettes (and have not stopped at any point) according to their longitudinal records up to the clinical visit when the adipose tissue biopsy was obtained. Ex-smokers were individuals who have successfully (and consistently) reported to have quit smoking cigarettes for at least three months prior to the adipose tissue biopsy. Non-smokers were individuals who never smoked according to the longitudinal questionnaire records. Other phenotypes such as age, body mass index (BMI), and alcohol consumption were also collected during the clinical visit. The alcohol consumption data were summarized as units per week, and then converted to grams/day (one unit of alcohol in the UK is defined as 7.9 grams (75)).

### Infinium HumanMethylation450 BeadChip data

The Infinium HumanMethylation450 BeadChip (Illumina Inc, San Diego, CA) was used to measure DNA methylation in both adipose and blood samples. Details of experimental approaches have been previously described (71, 76). At each CpG site, the methylation levels are characterized as a finite bounded quantitative trait ranging between 0 and 1, and represented as beta values. To correct for technical issues caused by the two Illumina probe types and two-color channels, the beta mixture quantile dilation (BMIQ) method (77) and background correction were performed for each sample. DNA methylation probes that mapped incorrectly or to multiple locations in the reference sequence were removed. Probes with more than 1% of subjects with detection P-value > 0.05 were also removed. All the probes are with non-missing values in blood samples and less than 1% missing subjects in adipose samples. Probes located on chromosomes X and Y were removed from the analysis. To check for sample swaps, we compared 65 single nucleotide polymorphism (SNP) markers on the array to genotypes for each subject, and removed subjects with incomparable genotypes. The methylation levels were normalized to N(0,1) prior to analysis.

### RNA-sequencing data

Twin adipose RNA-seq quality control and identification of batch effects have been previously discussed (78, 79). In brief, the sequenced paired-end reads (49 bp) were mapped to the human genome (GRCh37) by Burrows-Wheeler Aligner (BWA) software v0.5.9 (80), then genes were annotated as defined by protein coding in GENCODE v10 (81). Samples were excluded if they failed during library preparation or sequencing. Samples were only considered to have good quality if more than 10 million reads were sequenced and mapped to exons. The gene expression levels were quantified per gene, estimated as RPKM values (reads per kilobase of transcript per million mapped reads) and rank normal transformed prior to analysis. The genotype of each subject was used for identity checks in case of accidental sample swaps. After removing genes located on chromosomes X and Y, and non-coding transcripts, 17,399 genes were included in the gene expression analysis.

### Genotype data

Genotypes were available for all subjects in study. Genotyping of the larger TwinsUK dataset was performed using HumanHap300, HumanHap610Q, HumanHap1M Duo and HumanHap1.2M Duo 1M arrays. Imputation was done in two datasets separately, and subsequently merged with GTOOL. Genotype data were pre-phased using IMPUTE2 without a reference panel, then using the resulting haplotypes to perform fast imputation from 1000 Genome phase1 dataset (82, 83). We used 1000 Genomes Phase I (interim) as reference set, based on a sequence data freeze from 23 Nov 2010; the phased haplotypes were released Jun 2011. After imputation, SNPs were filtered at a MAF > 5%.

### Differential methylation and expression analyses

Principal component analysis (PCA) was used to identify potential batch effects. The association of smoking status with adipose methylation was examined using a linear mixed effect regression model (LMER) adjusting for batch effects (plate, position on the plate, bisulfite conversion levels, and bisulfite conversion efficiency), age, BMI, and alcohol consumption, family and zygosity structure. In blood samples, the association was tested adjusting for batch effects (plate and position on the plate), age, BMI, alcohol consumption, and 7 cell count estimations (plasma blast, CD8pCd28nCD45Ran, CD8 naïve, CD4T, NK, monocytes, and granulocytes), family and zygosity structure. The blood cell counts were calculated from the Horvath online calculator (84). A linear mixed effect regression model was applied as the data contained MZ and DZ twins. Family structure and zygosity were coded as random-effect terms, while all the other covariates were included as fixed-effect terms. Similarly, in the RNA-seq data analysis, the association of smoking status with expression levels was examined using LME adjusting for age, BMI, alcohol consumption (grams/day), GC mean, primer index, clinic visit date, family structure, and zygosity. Family structure, zygosity, primer index, and clinic visit date were taken as random-effect, and all the other covariates were included as fixed terms. For each CpG site, a full model that regressed all of the covariates was compared to a null model that excluded smoking status. The models were compared using the ANOVA F statistic. A genome-wide significance level was set at 1% false discovery rate for all analyses.

In order to account for mixtures of cell types in adipose tissue, we performed a EWAS using the reference-free approach proposed by Houseman et al (31). The method is similar to surrogate variable analysis (SVA) and independent surrogate variable analysis (ISVA), which is used to adjust for technical errors (e.g. batch effect) and confounders. In addition, the reference-free approach also includes a bootstrap step to account for the correlation in the structure of standard errors. Using this approach, we can estimate direct epigenetic effects that account for cell compositions and use bootstrap-based P-values to assess their significance. Due to the limitation that the reference-free approach can currently only be applied to datasets of unrelated individuals, we used 251 unrelated individuals from the original 542 twins and compared the top results between two EWASs.

To identify tissue-shared smoking differentially methylated signals across adipose and whole blood datasets, we compared the genome-wide FDR 1% signals across adipose and whole blood DNA methylation analyses. In whole blood samples we tested for association between smoking status and DNA methylation levels at 452,874 CpG sites in 86 current-and 481 non-smokers in blood. We compared the FDR 1% adipose DMS to 2,782 CpG sites that were associated with smoking in blood at FDR 1% (P = 1.14×10^−5^). We used a previously published lung cancer DNA methylation dataset (74) to further explore tissue-specificity at the 14 tissue-shared CpG-sites identified in both adipose and blood. We also checked smoking effects at the 14 tissue-shared CpG sites in 196 female subjects with skin tissue biopsies (33) applying a Bonferroni adjusted P-value of 3.6×10^−4^ as the significance threshold.

### ROC analysis

The sensitivity and specificity were calculated using receiver operative curve (ROC). The ROC analysis was performed using the pROC package (85) with the ‘lme’ function for logistic regression, where outcomes are categorized as smokers and non-smokers. We then used the ‘predict’ function to predict the expected probabilities under different combinations of predicting factors (methylation levels of 14 CpG sites and expression levels at 5 genes), and the ‘roc’ function to predict the sensitivity and specificity and draw the area under the curve. We selected 27 smokers and 145 non-smokers as a training set to construct a logistic model for smoking status classification, and then used the remaining set of 173 subjects (27 smokers) as a validation set, in which we obtained the AUC values. We repeated this procedure 1,000 times and report the average AUC values across 1,000 validation sets.

### Smoking cessation analyses

We quantified ‘reversal’ time by estimating the time (in smoking-quit years) required for ex-smokers to revert to 25% of the change in methylation towards non-smokers. For example, at cg05575921 in *AHRR*, the median level of methylation residual is −0. 234 in smokers and 0.037 in non-smokers, resulting in a 0.271 methylation change. Therefore, ex-smokers who reached methylation levels of −0.031, were classified as subjects who “reversed”. We quantified the proportion of subjects who reversed within different quit years. For example, at cg05575921, 6 ex-smokers quit for less than 1 year, but only one had methylation reverting to 25% of the methylation change towards non-smokers, therefore, the reversible rate is 16.7%. We quantified reversal at the gene expression level using the same approach.

### Methylation QTL (meQTL) analyses

Genome-wide meQTL analyses were performed testing for the association between common genetic variants and DNA methylation at CpG-sites in the 542 adipose tissue samples. We considered meQTLs at CpG-sites where at least one SNP was significantly associated with DNA methylation in *cis* (P = 5×10^−5^, as described in Grundberg et al. (71)), reporting the most significant SNP per CpG-site. In total, methylation levels of 102,461 CpG sites were associated with genetic factors in *cis*, and 25,531 sites in *trans*.

We tested the adipose meQTLs first by fitting a LME model regressed all the identified covariates, then performed a linear regression of the residuals on the SNPs using the MatrixeQTL R package (86). Results from meQTL analyses are presented at a P-value of 10^−5^ for the smoking-DMS, the smoking-DES, and at the smoking GWAS genetic variants. For meQTL analyses replicating the results from Loukola et al. (42) we applied a different threshold. Loukola et al. (42) conducted a genome wide association study of nicotine metabolite ratio, identifying many strongly associated SNPs in a 4.2Mb region on chromosome 19q 13. Among the 158 CpG sites within that region, 16 CpG sites showed statistically significant association with 173 SNPs. We compared our meQTLs findings to those from Loukola et al. (42) at a modified Bonferroni significance threshold of 1.81×10^−5^ (=0.05/16×173), and replicated meQTLs in 5 CpG sites (in *CYP2A7*, *ENGL2*, and *LTBP4* genes) (**S5 Table**).

### Direct comparison between methylation and gene expression levels

We compared the 542 subjects’ adipose methylation and gene expression levels at the five overlapping genes identified in the two genome-wide association analyses. Both the methylation and expression data were first adjusted for the covariates, and Spearman’s correlation test was then performed on the residuals.

### Metabolic disease risk phenotype analyses

We studied the impacts of smoking methylation signals on obesity and metabolic phenotypes. We explored 288 subjects (42 smokers and 246 nonsmokers) who had available DEXA profiles at or within up to 1 year of the adipose tissue biopsy. We compared the DNA methylation signals at the 42 smoking-DMS against adiposity phenotypes visceral fat mass, trunk fat, and android-to-gynoid fat ratio, adjusting for BMI. A significance level was set at a Bonferroni adjusted threshold of P= 5.7×10^−4^. We used a similar approach to test for phenotype associations with the 42 smoking-DES. To further investigate the effect of 42 smoking-DMS and 42 smoking-DES on weight gain after smoking cessation, the adiposity phenotype differences were obtained at two time-points (mean difference years = 5.1). We tested for correlations between the differences and methylation or expression levels at time point 1.

We used the R statistical software (https://www-r-project.org/) for all analyses and figures, and the regional plots were generated using the coMET package (87).

## Declarations

Ethical approval was granted by the National Research Ethics Service London-Westminster, the St Thomas’ Hospital Research Ethics Committee (EC04/015 and 07/H0802/84). All research participants have signed informed consent prior to taking part in any research activities.

## Data availability

Most of the datasets analysed in the current study are available under ArrayExpress accession number E-MTAB-1866 and EGA accession number EGAS00001000805 (adipose methylation and expression), GEO accession number GSE39279 (lung methylation (74)), and GEO accession number GSE90124 (skin methylation (33)). Additional individual-level data are not permitted to be shared or deposited due to the original consent given at the time of data collection. However, access to these genotype and phenotype data can be applied for through the TwinsUK data access committee. For information on access and how to apply http://www.twinsuk.ac.uk/data-access/submission-procedure/.

## Authors’ contributions

J.T.B. designed the study and outlined the main conceptual ideas. J.T.B., K.S.S., K.K., M.O., E.L., T.D.S., and K.H.P. supervised the work in each contributing research group. T.D.S., P.D., K.S.S, and J.T.B. generated the primary datasets. P-C.T. lead the data analysis. C.A.G., M.N.E., S.B., I.Y., J.E.C-F., T.H., T.C.M., A.V., M.M., K.W. and A.V. contributed data analysis. J.T.B. and P-C.T. wrote the article and all authors provided critical feedback and helped shape the research, analysis and manuscript. All authors read and approved the final manuscript.

## Supporting information

### Supplementary Figures

**S1 Figure.** Scatterplot of correlations between EWAS −log_10_P-values from the linear mixed effect model used in the current discovery study (y-axis) and results from Reference-free EWAS approach proposed by Houseman et al. (x-axis) (31).

**S2 Figure.** Smoking cessation and adipose DNA methylation profiles. DNA methylation levels at the 42 smoking-DMS and smoking status in 542 adipose samples. Subject groups include current smoker, subjects who quit smoking within one year, subjects who quit between 1 to 5 years, subjects who quit smoking more than 5 years, and subjects who never smoked. Fourteen CpG-sites located in genes with both smoking-DMS and smoking-DES are denoted with asterisks.

**S3 Figure.** Smoking cessation and adipose gene expression profiles. Gene expression levels at the 42 smoking-DES and smoking status in 542 adipose samples. Subject groups include current smoker, subjects who quit smoking within one year, subjects who quit between 1 to 5 years, subjects who quit smoking more than 5 years, and subjects who never smoked. Five genes with both smoking-DMS and smoking-DES are denoted with asterisks.

**S4 Figure.** Association between DNA methylation levels at the 42 smoking-DMS and future change in visceral fat mass (VFM) in 18 (red solid dots) and 228 subjects (grey hollow dots).

**S5 Figure.** Association between gene expression levels at the 42 smoking-DES and future change in visceral fat mass (VFM) in 18 (blue solid dots) and 228 subjects (grey hollow dots).

### Supplementary Tables

**S1 Table.** Four smoking-induced differentially methylated and expressed genes in blood samples.

**S2 Table.** Validation of the 14 smoking-DMS in 168 lung cancer tissues (74) and 195 skin tissues (33).

**S3 Table.** Replication of the 42 smoking-DMS in the LEAP cohort (38) with 104 smokers and non-smokers.

**S4 Table.** Previously-identified smoking genetic variants and their impacts on DNA methylation and gene expression in adipose tissue.

**S5 Table.** DNA methylation QTL (meQTLs) analyses at the chromosome 19 region from Loukola et al. (42), showing replication in TwinsUK adipose tissue samples.

**S6 Table.** Characteristics of TwinsUK and LEAP cohort (38).

**S7 Table.** Characteristics of 168 lung cancer (74) and 195 skin samples (33).

